# Waltz, AlphaFold and ESMfold Predictions on a Human Prion Protein Tiled Peptide Library Highlight a C-Terminal Region with Strong Conformational Sensitivity

**DOI:** 10.64898/2026.07.31.742043

**Authors:** Jennifer E. Grant, Sophia J. Vranicar

**Author notes:** LinkedIn: Jennifer Grant, Ph.D. | LinkedIn.

## Abstract

Short, linear sequence motifs within the human prion protein (PrP) may encode local aggregation tendencies that are not apparent from full-length sequence analysis. To map intrinsic amyloidogenic potential across PrP, we generated a complete one-residue-step library of overlapping 15-mer peptides from the 253-residue human PrP sequence and evaluated each peptide using WALTZ in both high-specificity and best-overall-performance modes, with full-length SNPeffect4/WALTZ output used for comparison. Peptide-level WALTZ analysis identified several candidate amyloidogenic regions, including an N-terminal signal-peptide segment spanning approximately residues 8–21/22 that was not detected in the publicly available full-length WALTZ/SNPeffect4 output. Predictions made using AlphaFold 3.0 also identified a short C-terminal area whose representative peptides formed either α-helix or pair of β-strands, indicating a region of potential higher susceptibility to conformational dynamics. This computational study supports the use of peptide tiling as a complementary screening strategy for identifying candidate short aggregation-prone motifs in PrP and other misfolding-associated proteins.

## Introduction

The human prion protein (PrP), encoded by PRNP, is a 253-amino-acid glycoprotein whose misfolding underlies a spectrum of fatal neurodegenerative disorders, including Creutzfeldt–Jakob disease, fatal familial insomnia, Gerstmann-Sträussler-Scheinker disease, kuru, and variant Creutzfeldt-Jakob disease [1–3]. Several physiological roles have been proposed for PrP, including contributions to neuronal development [4,5], synaptic plasticity [6], and metal homeostasis [7,8]. PrP contains an intrinsically disordered N-terminal region and a structured C-terminal domain, creating distinct sequence contexts in which local misfolding propensities may be difficult to detect from full-length analysis alone.

NMR [10,11] and cryo-EM structures [12,13], in tandem with traditional wet-lab proteomics and molecular biological approaches have provided much information about the misfolding of prions. Bioinformatics approaches including use of AI-based structural prediction approaches such as AlphaFold, ESMfold and Waltz have also added to this insight. However, our understanding of protein sequence determinants of prion misfolding is not sufficient enough to bring useful medications to patients. Realizing that PrP has disordered and structured regions, and that PrP misfolding within both types of regions can cause disease, we proposed that analysis of the virtual peptide library could reveal aggregation-prone motifs for prion misfolding.

To test whether short sequence segments encode aggregation-prone motifs, we generated a one-residue-step tiled library of 239 overlapping 15-mer peptides spanning the full human PrP sequence, following the sliding-window analysis strategy of Nystrom, *et al.*, to interrogate the aggregation tendency of 15-mer virtual peptides [14]. The 239 peptides were subjected to amyloidogenic potential screening using the Waltz predictive software [15, 16] and also secondary structure prediction using AlphaFold 3.0 [17, 18, 19] and ESMfold [20]. We have identified a predicted region of strong amyloid potential within the N-terminal signal sequence, and a second region in GPI anchor signal sequence that not only has strong amyloid potential but whose component amino acids are predicted to adopt either α-helix or β-strand conformation depending on the context of the peptide analyzed.

### N-terminal contributions to misfolding

Notably, in most NMR studies of prion proteins, the N-terminus remains disordered. The PrP N-terminus has attracted increasing attention as a potential modulator of misfolding. In particular, the signal peptide has been implicated in early events that influence PrP topology and aggregation. The N-terminal signal sequence has been previously identified in misfolding event. Specifically, synthetic peptides based on the prion N-terminus have demonstrated anti-amyloid activities [21]. Taking this one step forward into a study of prion accumulation, synthetic constructs containing hydrophobic signal-sequence segments fused to polycationic PrP motifs reduce proteinase K-resistant prion accumulation in infected cell models [22]. additional reports implicate the prion N-terminal signal sequence affects the degree to which the prion protein becomes properly inserted into endoplasmic reticulum [23]. Intriguingly, recent a recent study of a larger PrP N-terminal peptide PrP[1–110] implicates the human prion N-terminus in rapid neurodegeneration [24].

### C-terminal determinants of pathogenic conversion

The C-terminal domain of PrP (residues ∼125–228) forms the protein’s structured α-helical core and has been implicated in pathogenic conversion to the misfolded form PrP^Sc^. NMR studies of human PrP [10] and bovine PrP revealed the C-terminal globular domain where three α-helices arranged around a short β-sheet [11], where the C-terminal ends of helices α2 and α3 display local disorder. Disease-linked mutations such as E200K, D178N, and V210I cluster within this domain and destabilize its native conformation, promoting misfolding-competent intermediates [25–27]. In vitro fibrillization studies further demonstrate that isolated C-terminal fragments spontaneously adopt amyloid structures resembling PrP^Sc^ [28–30]. Although not the sole contributor to pathogenic conversion, destabilization of the C-terminal globular domain remains a central driver of PrP’s transition into self-templating conformations.

### Rationale for sliding-window peptide analysis

Full-length prediction tools emphasize global structural features, potentially masking shorter aggregation-prone segments. The role of short linear motifs in PrP local secondary-structure formation and amyloidogenicity is less well defined. Sliding-window peptide analysis provides a complementary strategy by isolating short sequence elements and evaluating their intrinsic structural propensities independent of the full-length fold. This approach revealed cryptic amyloidogenic motifs in the SARS-CoV-2 spike protein, where Nyström and Hammarström showed that short peptides can expose aggregation signals that are difficult to detect in the intact Covid spike protein [14].

Here, we similarly applied a systematic 15-mer sliding-window analysis to the human PrP sequence and evaluated each peptide using the WALTZ predictive algorithm [15] under both high-specificity and best-performance modes. Each peptide was also subjected to secondary-structure prediction using AlphaFold 3.0 [16–19] and ESMfold [20]. Our goal was to identify amyloidogenic motifs and contextualize them within the extensive literature on PrP biogenesis, membrane topology, and prion propagation. This peptide-tiling approach detected the N-terminal signal-sequence motif as amyloidogenic, and also identified an aggregation prone stretch of amino acids in the extreme C-terminus whose amino acids could participate in either an α-helix or a β-strand with equal ease.

## Results

### Global landscape of predicted amyloidogenicity (using Waltz)

A total of 239 overlapping 15-mer peptides were analyzed with WALTZ. The best-overall-performance mode identified a higher number of peptides and implicated more unique PrP residues than the high-specificity mode, consistent with its less stringent algorithm. Both peptide-level WALTZ modes implicated more unique residues than the full-length SNPeffect4/WALTZ comparison accessed at snpeffect.switchlab.org/uniprot/PRIO_HUMAN [31]. Importantly, the tiled-peptide analysis identified an N-terminal candidate amyloidogenic segment within the signal peptide that was not present in the full-length SNPeffect4/WALTZ output. As addressed below in our discussion, this N-terminal signal peptide area was previously identified as important in amyloidogenic events and an area of pharmacological intervention.

The overall results of performing the Waltz analysis using the best overall performance algorithm and the high specificity algorithm were largely similar and consistent with each other. Both analyses identified an N-terminal region and a C-terminal region likely to be amyloidogenic. A comparison of the location of different residues identified in Waltz Best Overall Performance mode and High Specificity mode is provided in Figure 1A and 1B, respectively. The N-terminal sequence identified by both algorithms was highly similar, representing residues within the PrP N-terminal signal sequence. Unsurprisingly, the best overall performance identified an additional two residues that the high specificity algorithm did not signal. In the C-terminal domain, both algorithms identified three swathes of primary structure as amyloidogenic. However, the high specificity algorithm identified significantly fewer residues (45 residues) than the best overall performance algorithm, which identified 76 residues.

**Figure 1.**
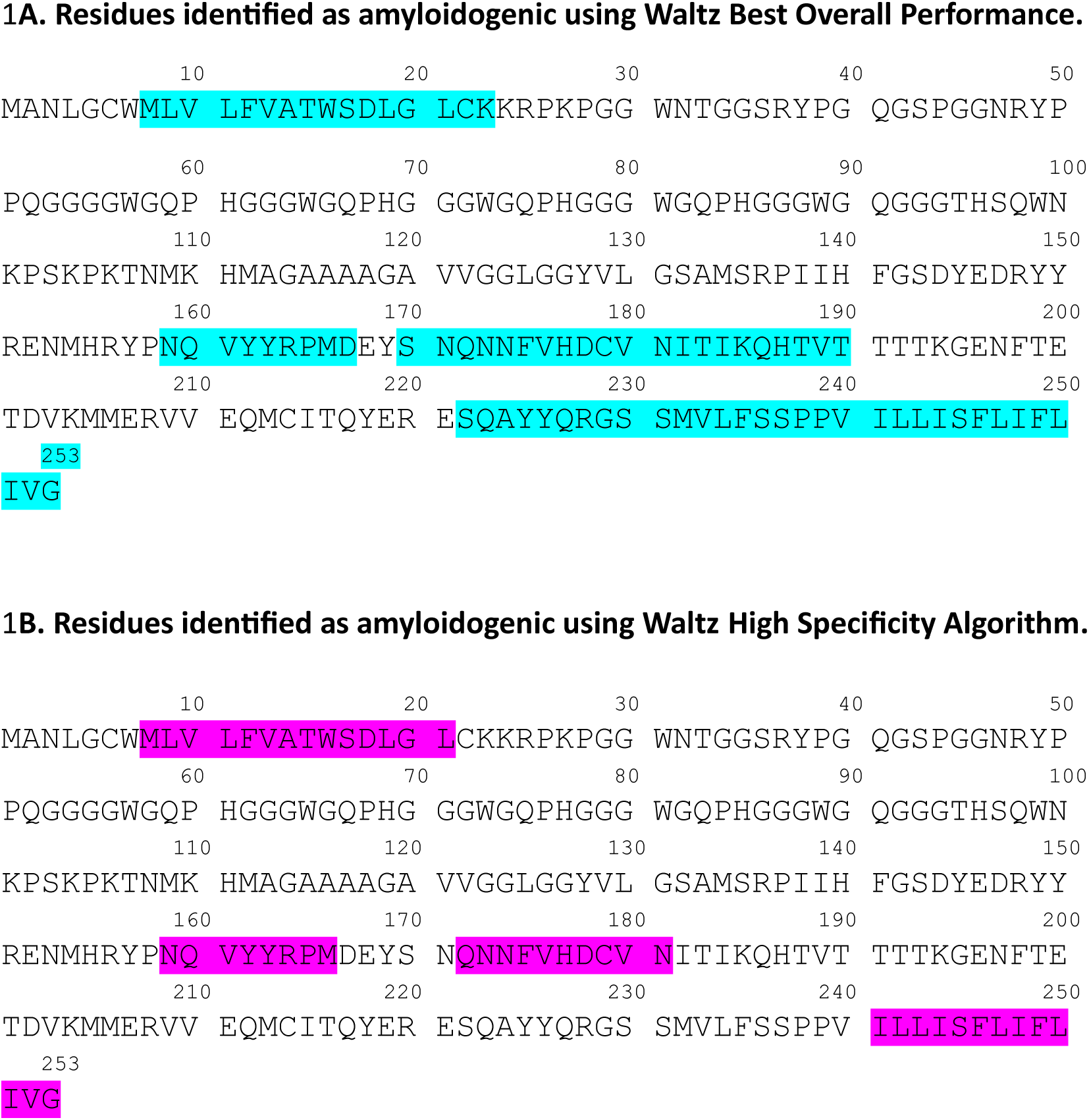
Waltz analysis for tiled 15-mer peptides. **1A.** Each peptide from the virtual library was folded with the Waltz Best Overall Performance Algorithm. **1B.** Each peptide from the virtual library was folded with the Waltz High Specificity Algorithm. Areas of red highlight indicates areas of predicted a-helix while underlining denotes areas where both a-helix and b-strand were predicted.

### WALTZ Identifies Predicted Multiple Amyloidogenic Regions

WALTZ predictions revealed several candidate amyloidogenic regions.

Use of the Best Overall Performance algorithm produced four candidate regions predicted to generate amyloidicity: PrP[8–22], PrP[159–167], PrP[169–190], and PrP[222–253]. As illustrated in Figure 1, most of these areas localize to the C-terminal globular domain of the prion protein. The exception was the sequence PrP[8–22] which resides in the extreme N-terminus of the prion protein and is largely associated with the prion signal sequence.

While use of the high-specificity algorithm reduced the number of amino acids identified, as expected, the results did faithfully capture the presence of four general regions: PrP[8–21], PrP[159–166], PrP[172–181], and PrP[241–253]. Again, one N-terminal sequence was identified, along with three C-terminal regions. Yet, while the PrP[8–21] and PrP[159–166] regions were identified largely intact, compared to the Best Overall Performance results, the high specificity algorithm markedly reduced the number of residues participating in the third and fourth regions, PrP[172–181], and PrP[241–253].

In Figure 2, the virtual peptides 213 to 231 are presented. As figure 2 demonstrates, one hundred percent of the nineteen virtual peptides were predicted to have amyloidogenic areas. The stretch representing [224–229], or AYYQRG, and the region [231–236], representing SMVLFS, were identified as amyloidogenic every time the entirety of the short sequence was present in a peptide.

**Figure 2.**
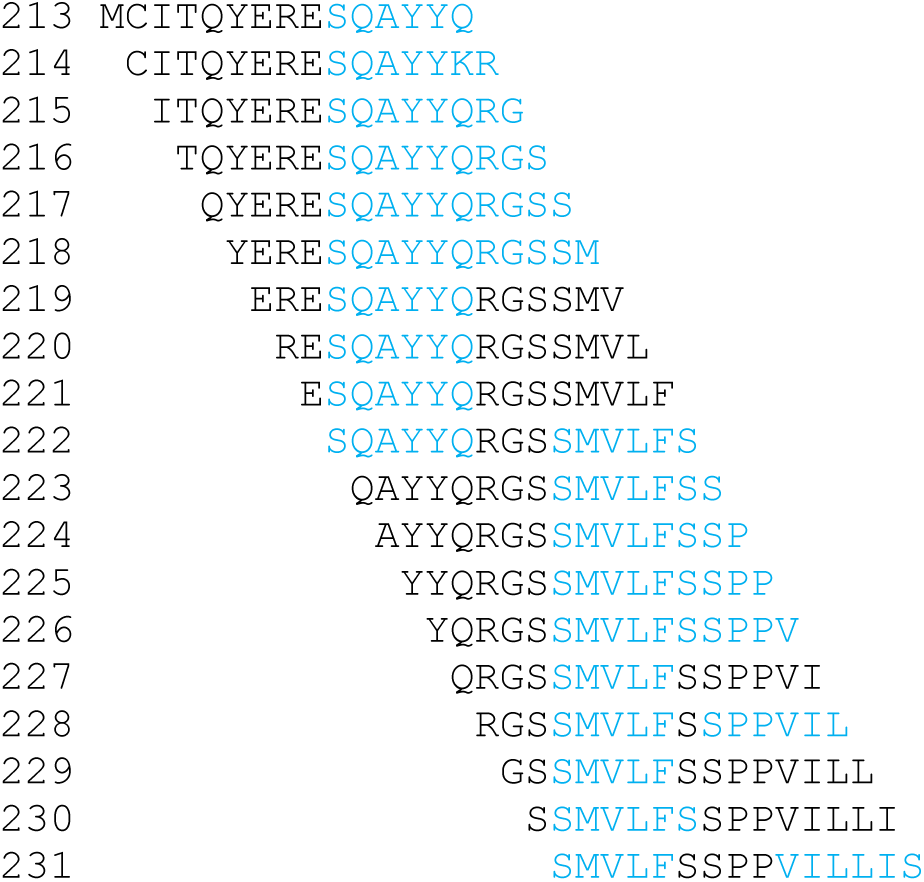
Waltz Best Overall Performance predictions for the major prion protein virtual library representing the region PrP[213–245]. Residues predicted to participate in amyloidogenic activity are highlighted in blue.

### Prediction of a N-terminal hotspot

Notably, tiled peptide-level analysis identified the amyloidogenic region PrP[8–21], a feature that was not identified in the SNPeffect4 database Waltz prediction set. This region was previously identified experimentally as a signal sequence that can participate in misfolding of the prion membrane protein.

### Prediction of Local Secondary Structure using AlphaFold

Secondary structure prediction employing AlphaFold 3.0/CoLab and using the same 15-mer peptide library identified seventy-one residues participating in α-helical content. The majority of these residues were located within the prion protein C-terminal portion (**Figure 3**). AlphaFold analysis of the peptide library failed to predict the presence of the β1 and β2 strands presented in the NMR structure 1QLM [10].

**Figure 3.**
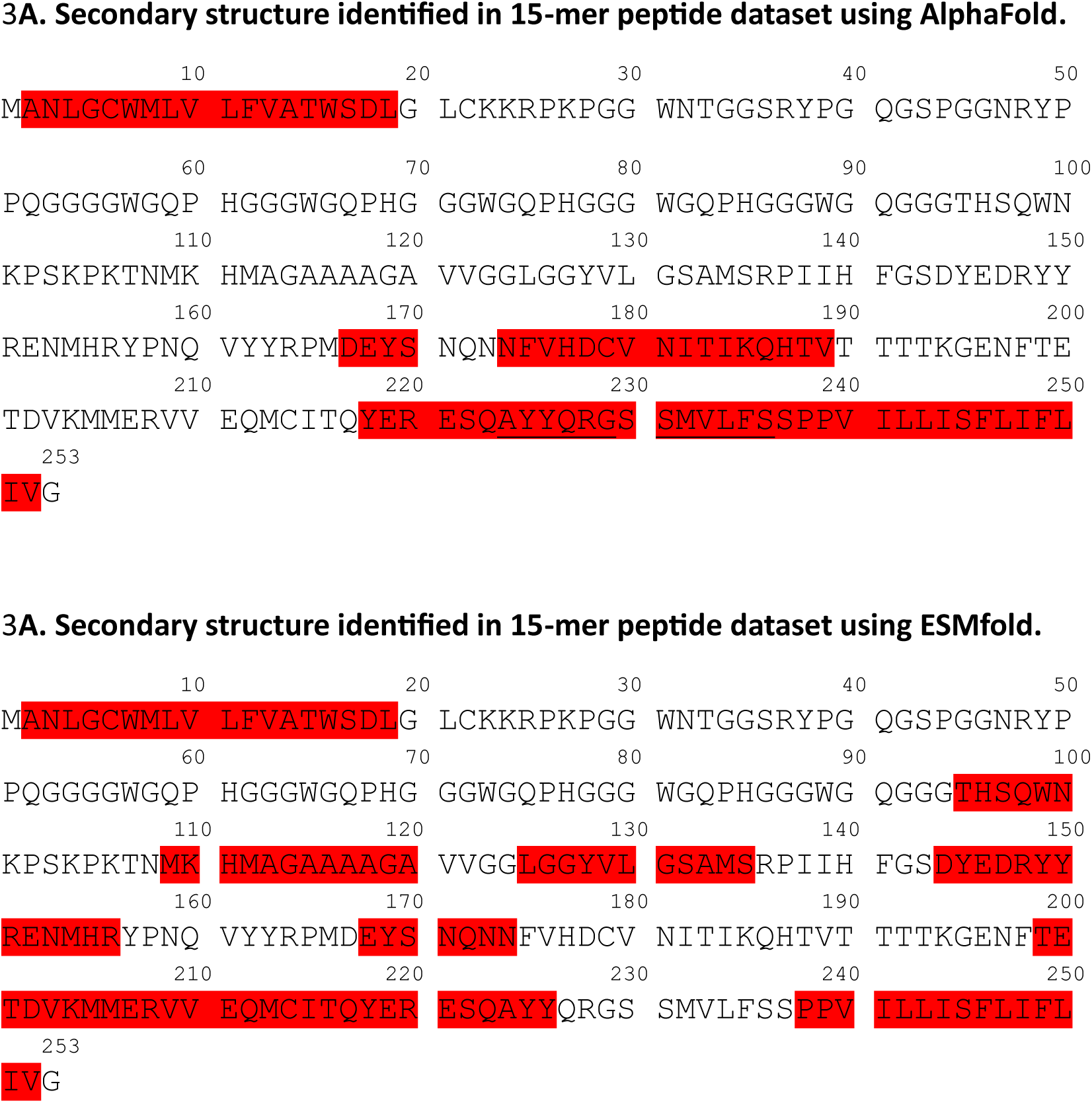
Secondary structure as revealed by predictions using tiled 15-mer analysis. **3A.** Each peptide from the virtual library was folded with AlphaFold 3.0. **3B.** Each peptide from the virtual library was folded with ESMfold. Areas of red highlight indicates areas of predicted α-helix while underlining denotes areas where both α-helix and β-strand were predicted.

Curiously, AlphaFold identified a separate C-terminal region containing twelve residues that participated in β-strands. These β-strand predictions occurred in peptides that incorporated the positions PrP[224–229] and PrP[231–236]. However, whether these residues participated in alpha helix or β-strand depended on which 15-mer was being folded (**Figure 4**). Only four peptides out of the entire virtual library predicted formation of β-strands, and the percent of β strand content depended strongly on the ability of the peptide to a pair of β-strands that presumably interact to form a β-sheet feature.

**Figure 4.**
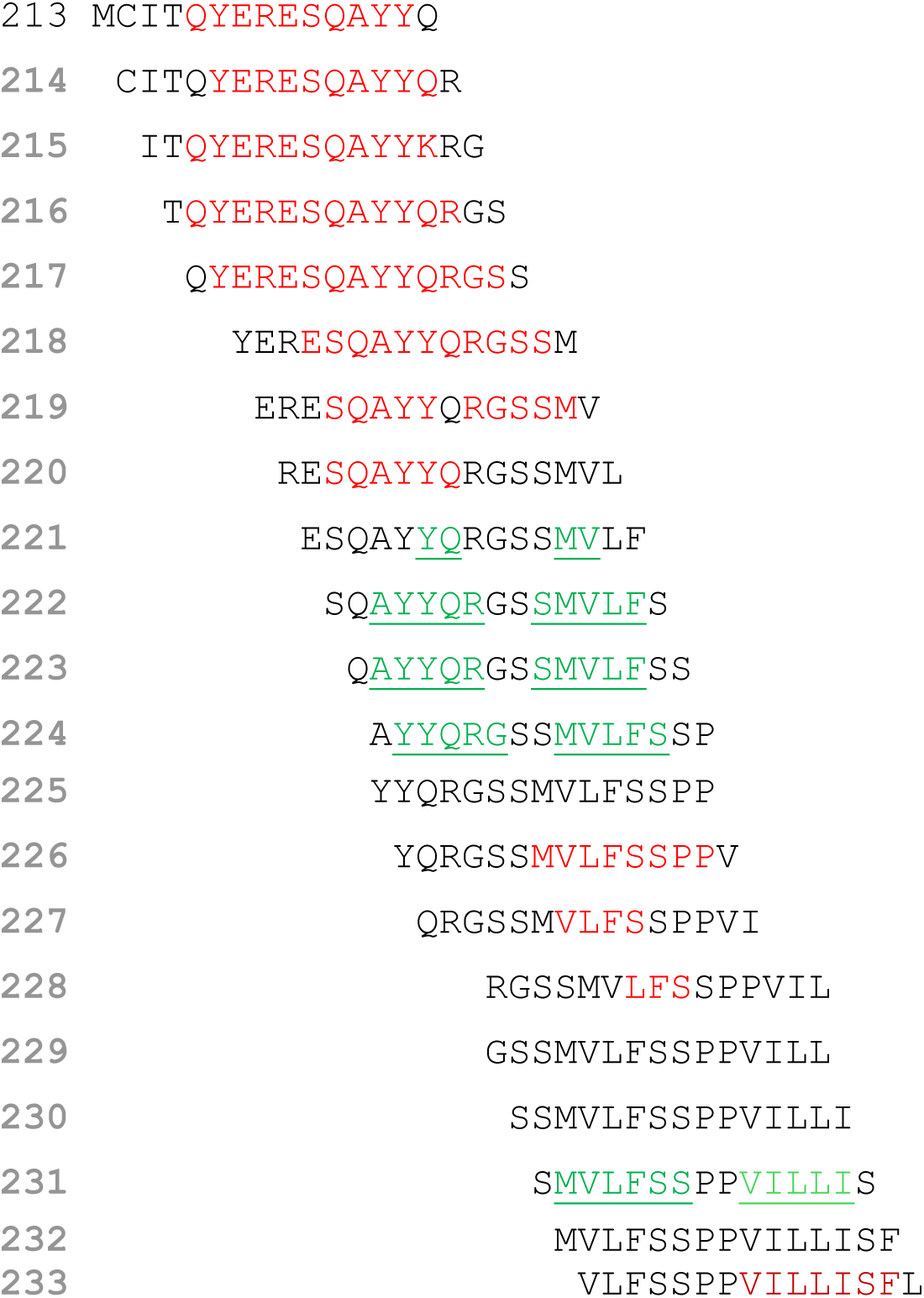
AlphaFold secondary structure predictions for the major prion protein over the region PrP[213–245]. For each peptide, residues predicted to participate in an α-helix are highlighted in red while residues participating in β-strand are highlighted in green and underlined.

### Prediction of Local Secondary Structure using ESMfold

The same 15-mer peptide library was analyzed for predicted secondary-structure content using AlphaFold 3 and ESMfold peptide models. ESMfold predictions of α-helix content within the 293 tiled peptides identified 105 residues participating in α-helical content, which closely recapitulated the AlphaFold Database prediction for human full-length PrP of 110 residues. ESMfold analysis of the peptide library failed to predict the presence of any β-strands, not even the β1 and β2 strands presented in the NMR structure 1QLM [10].

## Discussion

### Overall Disposition of Predictions

Within the two Waltz datasets, only four regions were predicted to have areas of high aggregation potential. One discrete region spanning PrP[8–22] was identified in the human prion N-terminus. This region is a unique identification not elucidated in the SNPeffect4 dataset that was previously published online. The additional three regions identified as having strong amyloid potential are in the C-terminal region, a region where there are a surfeit of experimental 3D structures and a deep literature base for understanding the impact of point mutations on PrP structure and function.

Notably, over the 478 15-mer peptide predictions made using either AlphaFold 3.0 or ESMfold, analysis of only one region in PrP contained residues that in some peptides participated in both β-helix and β-strand secondary structure. These chameleonic residues, identified only in the AlphaFold 3.0 dataset, spanned the region PrP[224 to 229] and PrP[231–236], spanning only ten residues.

The divergence between AlphaFold/ESMFold secondary-structure assignments and WALTZ amyloidogenicity predictions emphasizes that these tools measure different properties. AlphaFold and ESMFold provide structure predictions optimized primarily for folded proteins or protein assemblies, whereas WALTZ identifies short sequence segments with amyloid-forming propensity based on position-specific scoring. Therefore, a lack of predicted β-strand in an isolated model does not rule out amyloidogenic behavior, and a WALTZ-positive result does not by itself establish experimentally realized amyloid formation.

### The Prion N-terminus

Sliding-window peptide analysis identified candidate amyloidogenic PrP motifs that are not apparent in full-length WALTZ/SNPeffect4 output. That the N-terminal signal peptide was identified is notable. The PrP signal sequence influences PrP biogenesis and topology and has been linked experimentally to the generation of disease-associated transmembrane PrP forms. The importance of the N-terminus has been documented in the literature, and Waltz analysis of the tiled peptide library correctly identified this region as amyloidogenic.

Furthermore, PrP-derived peptide constructs containing hydrophobic signal-sequence motifs and polycationic PrP-derived segments have been reported to reduce proteinase K-resistant prion accumulation in infected cell cultures, and related cell-penetrating peptide constructs have been reviewed for anti-prion and anti-amyloid effects. While these experimental findings do not prove that residues 8–21/22 are amyloidogenic in the native PrP precursor, they support the broader idea that PrP N-terminal signal-sequence-derived peptides can have measurable biological and biophysical activities. The Waltz predictions identified here suggest that aggregation potential may be a factor.

Overall, these findings support peptide tiling as a useful hypothesis-generating complement to full-length prediction. The approach highlights several candidate aggregation-prone PrP motifs, including a tiled-peptide-only N-terminal signal-peptide segment, which can now be prioritized for orthogonal computational analysis and experimental validation. The most important next step is to evaluate whether the predicted N-terminal peptide forms amyloid-like assemblies in vitro and whether signal-sequence variants alter PrP biogenesis or aggregation-related phenotypes in cellular models.

### The Prion C-Terminus

In the human prion protein, residues 125-228 are considered part of the structured region of the C-terminus that includes the globular domain. This globular region has long been of interest in pathogenic amyloid formation. Much attention has been given to the α2, α2/α3 loop, and α3 helix. Within this globular domain, two prominent helices, the α2 and α3 helices, are known to impact protein stability.

In this work, the Waltz data using the best overall performance algorithm identified that, starting at residue 167, sixty-two residues were identified as having amyloidogenic potential. Much of this content includes the α2 and α3 helix. This finding further highlights the prion C-terminus’ role in amyloidogenic events.

When AlphaFold3 was used to predict regions of α-helical content within the entirety of the prion, fifty-five unique residues were identified as likely to participate in α-helix. In contrast, AlphaFold 3.0 analysis of the tiled peptide library assigned β-strand content to only ten unique residues in the C-terminal portion of PrP, whereas no β-strand assignments were observed in the N-terminal or central regions. Interestingly, these twelve residues were found also in peptides predicted to adopt α-helix structure, depending on which the exact 15mer peptide for which the prediction was made. Of the two stretches of amino acids, PrP[224–229] resides within the α3 helix of the 1QLZ NMR structure, while the amino acids PrP[231–235] are located distal to the α3 helix.

Many of the residues within PrP[224–229] and PrP[231–236] could reside in either alpha helix or β-strand conformation, depending on which 15mer was being evaluated, according to the AlphaFold predictions (**Figure 4**). This may indicate a region particularly susceptible to conformational change. Furthermore, PrP[224–229] lies at the extreme end of the α3 helix, which is where the ordered region of the 1QLZ NMR structure stops. PrP[231–236], on the other hand, lies distal to the α3 helix. It is particularly intriguing to consider whether these two regions, which lie adjacent to each other and are separated by only two residues, could adopt two β-strands that then interact to create a β-sheet structure. Indeed, Gong, *et al.,* [32] demonstrated differences in the solvent accessibility between a recombinant protein fragment and a scrapie model of the Syrian hamster, where the solvent accessibility differential was most pronounced within the extreme C-terminus of the prion protein corresponding to the region represented in the human prion by residues PrP[224–229].

### C-terminal Prion Mutations

The prion C-terminus harbors a signal sequence for a glycophosphoinositol (GPI) anchor, which direct the prion protein, and other similarly tagged proteins, to lipid rafts that facilitate signaling interactions [33]. The GPI anchor is also implicated in the proteolytic processing and shedding of prions. Within PrP[224–229] lies Y226. A patient with a premature stop codon at position 226, noted as the Y226X mutation, experienced a particularly severe case of prion disorder that was associated with formation of an unglycosylated 7kDa prion fragment [34]. The lack of a glycosylphosphatidylinositol anchor was identified as the signal reason for predisposition to amyloid plaque formation, emphasizing the importance of this residue.

Within the sequence PrP[231–236] lies 232M, a second mutation that also interferes with the GPI anchor. Work to understand how the mutation M232R impacts patients [35] revealed that this region participates is necessary for the prion to receive the GPI anchor required for proper translocation into the endoplasmic reticulum. The full sequence of the prion C-terminal signal sequence is PrP[231–253], which overlaps with PrP[231–236], half of the residues that the peptide tiling strategy identified as being able to participate in either α-helix or β-strand.

The peptide tiling approach revealed that residues within the two stretches PrP[224–229] and PrP[231–236] are positioned within a context-dependent environment, and uniquely positioned affect to GPI anchoring, lipid raft positioning, and similarly proteolysis and shedding processes. It is tempting to speculate that formation of β strands within both PrP[224–229] and PrP[231–236] could result in a limited β-strand structure that interferes with normal prion biology, resulting in misfolding events that ultimately result in prion disease.

### Limitations

This study has several limitations. First, WALTZ predictions are computational and may be influenced by hydrophobicity, peptide length, and signal-peptide composition; therefore, the N-terminal hit should be considered a candidate motif rather than a demonstrated amyloid-forming segment. Second, the 15-mer window length was chosen to capture short linear motifs while maintaining enough sequence context for peptide-level prediction, but the robustness of the result should be evaluated with additional peptide sizes. Third, overlapping peptide calls create a non-independent testing structure, and the false-positive burden should be assessed with orthogonal predictors and appropriate control sequences, such as unrelated human signal peptides. Fourth, AlphaFold 3 and ESMFold are not optimized for experimentally validating isolated short-peptide conformations, so their outputs are best used as qualitative structural annotations. Finally, biological relevance will require experimental validation, including peptide aggregation assays, structural characterization, and cellular tests of signal-sequence processing or topology.

### Future Directions

This work focused on a comparison of predictions on a virtual library of 15mer peptides using the WALTZ best overall performance algorithm, the WALTZ high specificity algorithm, AlphaFold 3.0, and ESMfold. It might be helpful to expand the analysis using the same peptide library to include additional amyloid or aggregation predictors such as AGGRESCAN, TANGO, PASTA, or Ribbon Fold.

A second and perhaps more intriguing line of inquiry would be to establish whether performing this analysis using virtual peptide libraries reflecting pathogenic mutations may reveal how single-residue substitutions reshape local aggregation potential.

### Summary

While use of the WALTZ algorithms to understand the aggregation potential of regions of the prion protein highlighted the aggregation of an N-terminal signal sequence known to be involved in misfolding events, and AlphaFold 3.0 predictions emphasized ten residues in or near the α3 helix that were predicted to participate in both β-helical and β-strand content, use of ESMfold did little to add to our ability to predict sensitivity of conformation within the prion protein.

These prediction data on the human prion protein, using a 15mer virtual peptide library, correctly identified the N-terminal region of the prion protein as an area of interest in prion aggregation biology, highlighting the potential contribution of AI-based software such as WALTZ and other folding prediction software in understanding protein aggregation in proteins that are less well-characterized.

## Methods

### Peptide Tiling

The analysis used the human PrP sequence from UniProtKB [36] accession Q53YK7 (major prion protein), which corresponds to the 253-amino-acid precursor protein, except that the exact input sequence utilized V129 sequence variant, as opposed to the M129 canonical sequence. A virtual peptide library was generated from the full precursor sequence using a one-residue sliding window and a fixed peptide length of 15 amino acids. For a 253-residue input sequence, this procedure yields 239 peptides with coordinates 1–15, 2–16, 3–17, and so forth through 239–253. Only complete 15-mer windows were retained.

### Amyloidogenicity Prediction

Each 15-mer peptide was analyzed with WALTZ (waltz.switchlab.org), a position-specific amyloid-propensity prediction method developed to distinguish ordered amyloid-forming sequences from amorphous β-aggregating segments. Two WALTZ settings were applied: high-specificity mode, which prioritizes lower false-positive rates, and best-overall-performance mode, which balances sensitivity and specificity. A peptide was counted as positive if WALTZ returned an amyloidogenic segment within that 15-mer under the specified mode. Residue-level counts were calculated by marking each PrP residue covered by at least one positive peptide and counting each residue once, regardless of the number of overlapping positive peptides that included it.

Full-length PrP amyloidogenicity was assessed using the SNPeffect4 WALTZ annotation for UniProtKB P04156 [31] to provide context for the tiled-peptide analysis.

### Secondary-Structure Prediction

Secondary-structure tendencies were evaluated for each 15-mer using AlphaFold 3 and ESMFold. AlphaFold 3 predictions, obtained through the fall of 2025, were generated for each peptide as single-chain peptide inputs through use of ChimeraX software (reference here). ESMFold predictions, obtained during the spring of 2026, were generated by deploying a python script that made requests through the ESMFold API. In all cases, predicted structures were saved as PDB files and analyzed in PyMOL 31.8 using its secondary-structure assignment algorithm. Residues assigned to β-strand or α-helix were counted once at the if they appeared in any peptide model with the corresponding assignment.

For full-length structural comparison, the AlphaFold Database model AF-P04156-F1-v6 was used obtain its prediction for full-length human PrP. A full-length ESMFold model of the same sequence was also generated or retrieved for comparison. These full-length models were used to compare the location of predicted secondary-structure elements with the peptide-level results, not as direct evidence of amyloid formation.

**Table 1.**
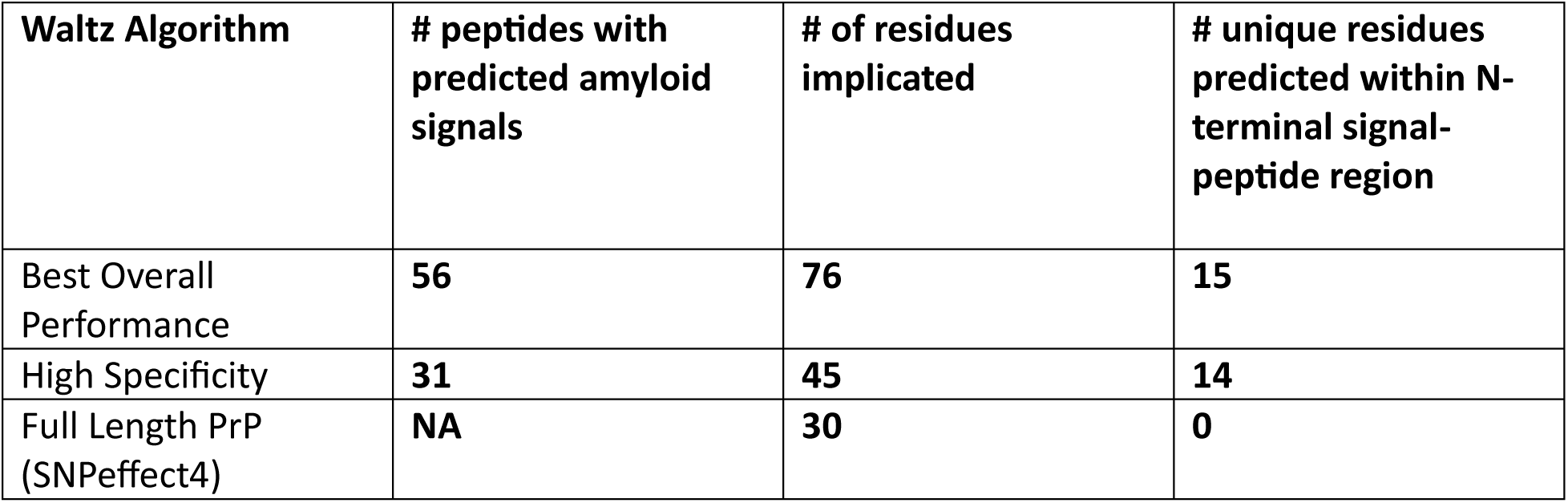
WALTZ-based amyloidogenicity predictions for tiled PrP 15-mers and full-length PrP.

**Table 2.** Secondary-structure assignments from peptide-level and full-length structure predictions.

| | # peptides with predicted β-strand content | # residues predicted within β-strand | # peptides with predicted α-helix content | # residues predicted in α-helix |
| --- | --- | --- | --- | --- |
| Peptide scan, AlphaFold | 5 | 12 | 34 | 71 |
| Full length PrP, AlphaFold | NA | 5 | NA | 110 |
| Peptide scan, ESMfold | 0 | 0 | 68 | 105 |
| Full length PrP, ESMfold | NA | 0 | NA | 111 |

## Acknowledgements

This work was supported by the Andrew G. Schneider Professorship to J.E.G.

## Declaration of AI Use

AI software AlphaFold 3.0, Waltz, and ESMfold were used to generate predictive data. CoPilot was used to strengthen the grammar of the manuscript.

## Author Contributions statement

JEG was responsible for conception and design of this project, drafting of the paper, and the final approval of the version to be published. Both authors contributed to critical revision of the intellectual content of this work and agree to be accountable for all aspects of the work.

## Data Availability Statement

Individual predicted poses are available upon request; this is a large dataset that was curated by hand. Compiled outcomes of the prediction set representing analysis of the virtual library are available through Minds@UW.

